# Spatio-temporal analysis identifies hotspots of marine mammal strandings along the Indian coastline: implications for developing a National Marine Mammal Stranding Response and Management policy

**DOI:** 10.1101/2021.01.20.427409

**Authors:** Sohini Dudhat, Anant Pande, Aditi Nair, Indranil Mondal, Kuppusamy Sivakumar

**Affiliations:** Wildlife Institute of India, Chandrabani, Dehradun – 248001, Uttarakhand, India

**Keywords:** Emerging Hotspot Analysis, geospatial tool, marine mammal occurrence, marine policy

## Abstract

Marine mammal strandings provide vital information on their life histories, population health and status of marine ecosystems. Opportunistic reporting of strandings serve as a potent low-cost tool for conservation monitoring of these highly mobile species. We present the results of spatio-temporal analyses of marine mammal stranding events to identify hotpots along Indian coastline. We collated data over a long-time frame (~270 years) available from various open access databases, reports and publications. Given the inadequacy in data collection over these years, we grouped data into four major groups viz. baleen whales, toothed whales, small cetaceans and dugongs. Further, we described the trends in data for marine mammal sightings, incidental mortalities, induced mortalities and stranding events using the last group for spatio-temporal analysis. Annual strandings along the Indian coast has increased considerably in the recent years (11.25 ± 9.10 strandings/ year), peaking in the last two years (2015-17, mean = 27.66±12.03 strandings /year). We found that number of strandings spiked in June- September along the west coast and December- January along the east coast. We identified several sections of coastline which have consistently received comparatively higher number of stranded animals (0.38 - 1.82 strandings/km) throughout the study period. Use of novel geospatial tool ‘Emerging Hotspot Analysis’ revealed new and consecutive hotspots along the north-west coast, and sporadic hotspots along the south-east coast. Despite the challenges of working with an opportunistic database, this study highlights critical areas to be prioritized for monitoring marine mammal strandings in the country. We recommend establishing regional marine mammal stranding response centres at the identified hotspots coordinated by a National Stranding Monitoring Centre with adequate funding support. Regular conduct of stranding response programs for field veterinarians, frontline personnel focused around identified stranding hotspots would help develop a comprehensive picture of marine mammal populations in Indian waters.

## 1. INTRODUCTION

Understanding marine mammal stranding patterns serves as a potent and inexpensive tool to study their populations around the world (Leeney et al. 2008; Mullins, 2008; Pikesley et al. 2012; Foord et al. 2019). Marine mammal strandings provide crucial information on their life histories (Arrigoni et al. 2011), population sizes (Prado et al. 2016), impact of anthropogenic activities on marine ecosystems as well as overall health of oceans (Prado et al. 2016; Coombs et al. 2019). Marine mammal strandings are known to occur due to various biological (diseases and parasitism-(Gonzales-Viera et al. 2011; Cuvertoret-Sanz et al. 2020); hearing impairments hindering navigation - Mann et al. 2010); environmental (unusual tides, electrical storms - Walker et al. 2005; cyclones- (Rosel and Watts 2008); geomagnetic anomalies, echolocation distortion Sundaram et al. 2006 and; anthropogenic factors (vessel strikes - Laist et al. 2001; net entanglement - Arbelo et al. 2013; Cassoff et al. 2011; Vishnyakova and Gol’din 2015; noise generated by dredging, oil drilling, naval exercises for SONARs -(Romano et al. 2004; Taylor et al. 2004; Weilgart 2007) and; marine pollution- (Secchi and Zarzur 1999; Law et al. 2012). Predicting marine mammal strandings though is often difficult due to various ecological factors at play to determine the location of stranding. However, opportunistically available marine mammal strandings offer a crucial tool to identify marine mammal distribution (Maldini et al., 2005) as a relatively cheaper monitoring method over dedicated primary surveys (Betty et al., 2019).

The foraging areas of marine mammals often overlap with fisheries as both utilize the most productive zones in the ocean (Goetz et al. 2015; Weinstein et al. 2017). This overlap often results in accidental net entanglement (Reeves et al. 2013) or injuries/ mortalities due to hooking in longlines (Gilman et al. 2006). Movement of high speed vessels and ships near port and other areas also pose a risk of collision with marine mammals as they surface to breathe (Groom et al. 2004; Constantine et al. 2015). Noise pollution in the marine environment can interfere with the echolocation of marine mammals, causing damage to the auditory system or lead to ‘auditory masking’ (i.e. diminishing the ability of the animal to detect a relevant sound) (Engel et al. 2004; Compton et al. 2008). Besides accidental mortalities or disorientation due to human-mediated factors, coastal topography can also play a major role in clustering stranding events along certain sections of coastlines (Sundaram et al. 2006). Factors such as gently sloping beaches, also known as ‘acoustic dead zones,’ distort acoustic signals of cetaceans,, confounding orientation and leading to stranding (Sundaram et al. 2006).

Physical or environmental factors such as near shore surface currents, local wind patterns (Norman et al. 2004) and coastal topography (Brabyn and McLean 1992; Hart et al. 2006; Leeney et al. 2008) play a huge role in bring the carcasses of dead animals to the shore. Further, temporal variations in animal movements (e.g. migrating to warmer waters for breeding; Jenner et al. 2001) as well as changes in intensity of fishing, tourism and other human activities might result in deaths due to boat strikes and entanglement (Sutaria et al. 2015; Vishnyakova and Gol’din 2015) which could explain stranding patterns in certain seasons at certain sections of the coastline.

In India, an extensive coastline (~8000 km) and large Exclusive Economic Zone (area 2,305,143 Km^2^) makes it difficult for monitoring agencies to manage marine mammal strandings. Further, Indian waters support marine mammals (~ 34 species MMRCNI, http://www.marinemammals.in/) utilizing various oceanic zones, with varying anthropogenic and natural threats, creating complex challenges for their management. Conservation and management of marine mammals over the Indian coastline remains a major challenge. Therefore, identifying patterns in marine mammal strandings are crucial to direct efforts to regions with marine mammal populations and help the agencies such as State Forest Departments for developing effective management interventions. Given this context, in the present study, a) we evaluate the general trends in marine mammal occurence; b) analyse temporal and spatial patterns associated with marine mammal strandings; c) and identify spatio-temporal hot-spots for marine mammal stranding events along the Indian coastline. We consider these results to broadly apply for formulating an effective National Marine Mammal Stranding Response and Management policy in India.

## 2. METHODS

In this study, we compiled data on marine mammal strandings in India from various published and unpublished sources and organised them into a single dataset. The data sourcing, compilation and analysis was conducted using following steps:

### 2.1. Data sources

Data on marine mammal records from Indian coast was accessed from peer-reviewed publications, grey literature such as unpublished reports, thesis, newspaper articles and open access database. Data other than strandings (sightings, accidental net entanglements and hunting) was also collected to understand the species composition of marine mammals along Indian coastlines and to give a comparative account of number of cases in each of these categories.

We organised the data available through a period of 269 years (1748 to 2017) from multiple sources as listed below:

2.1.1. Publications- We extracted information pertaining to marine mammal from over 90 publications and one doctoral thesis. We found 440 records which included strandings (n=225), sightings (n=70), accidental net entanglements (n=139) and hunting (n=6) incidences.

2.1.2. Marine Mammal Research and Conservation Network of India (MMRCNI) database (http://www.marinemammals.in/; accessed on 6^th^ August 2018): This is an open access online database and virtual networking platform which receives voluntary submissions of marine mammal sighting and stranding information by researchers, students as well as the general public. We acquired 486 records from this source, which includes sightings (n=102), accidental net entanglements (n=80), hunting records (n=17) and strandings (n=287) of marine mammals.

2.1.3. Technical report on ‘Assessment of dugong distribution, habitat and risks due to fisheries and other anthropogenic related activities in India’ (Sivakumar and Nair, 2013): We accessed this institutional dataset on dugong sightings and strandings obtained using a standardised dugong catch/ bycatch questionnaire developed by the Convention on Migratory Species – United Nations Environment Program Dugong Memorandum of Understanding. The data acquired included sightings (n=461), accidental net entanglements (n=19) and hunting records (n= 34) and strandings (n=108) of dugongs.

2.1.4. Dataset of the project ‘Recovery of Dugongs in India and their habitat: an integrated participatory approach’ (Nair and Sivakumar 2013; Srinivas et al. 2020)). Under this project, data on dugong distribution is being collected from Gulf of Kutch (Gujarat), Palk Bay & Gulf of Mannar (Tamil Nadu) and Andaman & Nicobar Islands through an informant network comprising of fishers, SCUBA divers, frontline personnel from agencies such as State Forest Departments, Indian Coast Guard, Indian Navy and Marine Police. We extracted records for marine mammal sighting (n= 19), net entanglement (n=3) and stranding (n=46) from this dataset.

2.1.5. Local newspaper articles reporting strandings: Around 47 reports of marine mammals washed ashore were gathered from recent (articles from 2008 to 2016) regional language newspapers of Tamil Nadu (n=32), Pondicherry (n=2), Andhra Pradesh (n=3), Kerala (n=5) and Karnataka (n=5).

### 2.2 Data compilation

We compiled data from all these sources into a spreadsheet. Information about each stranding/ sighting incidence was compiled without any changes as provided by the author/ expert/ informant. We included information about the event (live stranding/ dead stranding/sighting), species name (if identified), date, location, and additional information if available (condition of carcass, age class, sex and any other observations of the author/expert/ informant regarding the incident) in the spreadsheet. We eliminated the records falling outside of Indian territorial waters (from Sri Lanka and Pakistan, received through MMRCNI database) as they exceed our range of study. MMRCNI database comprises of information from publications as well as from informants (locals and researchers). As our dataset also consisted of records from publications, some records were found to be overlapping (n=168) and were eliminated to avoid repetition.

The coordinates of each location in the dataset were overlaid in Google Earth Pro version 7.3 for visualization. Coordinates for few locations (n = 58) were either not available or found to be placed too far inland from the mentioned site. These locations were rechecked with the database and coordinates of the nearest shore of the mentioned village/ district was added manually, by noting the coordinates of approximate midpoint of the coastline.

### 2.3 Data processing

The problem of misidentification of cetacean species in the reports and publications have been highlighted in the earlier reviews on cetacean research in India (Kumaran 2002). To avoid problems due to this, we pooled all the records into four taxonomic groups i.e. baleen whales, toothed whales, smaller cetaceans (dolphins and finless porpoise) and dugongs to look at spatio-temporal patterns as a group rather than for each species considering biases arising from these misidentifications.

The terms used for reporting the type of marine mammal records was loosely used in the sourced information, for example, washed ashore, floating etc. Thus, we reclassified the information and grouped them under following heads to ease our process of analysis:

a. Strandings: animals found washed ashore dead or floating near shore or found stranded alive and attempted to being rescued.
b. Sightings: direct sighting of live animals
c. Incidental mortality: animals captured in gill nets, other net entanglements, boat strikes.
d. Induced mortality: animals reported hunted, killed, or driven ashore.
e. Not known: the state of the animal was not clearly mentioned by sources.

Further, the data was classified on the location of each record with respect to east or west coast of India. Records from the island archipelagos of Lakshadweep and Andaman & Nicobar Islands were classified as islands records.

### 2.4 Data Analysis

The stranding records compiled above (n=632) were further used for temporal and spatial analysis.

#### 2.4.1 Temporal analysis

The stranding reports before 1975 were sporadic (n =207 records spread over 227 years, from −1748 to 1975) and thus were excluded from further analysis.

Initially, we investigated the changes in numbers of strandings across decades collectively for all groups. Later, to understand the differences in stranding numbers on a finer scale, we calculated the rate of strandings of each group by dividing the coastline into 50 Km sections using ArcGIS 10.5 (procedure mentioned in detail in the spatial analysis section below) and assigned the total number of strandings to these sections. We obtained per kilometre stranding rates for each group for each 50 km section. The stranding rates for each section in the coastline was then pooled to compare the stranding rates of each group across both the coastlines.

These rates were further used to visualise yearly and monthly patterns for each group of marine mammals separately. We used the Mann Whitney U test to assess the significance of differences in the stranding rates of each group across the two coastlines. The discontinuous coastlines of islands made it difficult to be divided into section by the same method, the rates of strandings could not be calculated for the islands. Thus, the islands were excluded from temporal analysis.

#### 2.4.2 Spatial analysis

We divided the coastline of mainland India into segments of 50 km each using ‘Split’ tool in ArcGIS 10.5. A total of 170 segments were created and each stranding location along the coasts were assigned to the nearest segment using the ‘Near Analysis’ feature in ArcGIS 10.5. Further, stranding rates per km (*s_r_*) were calculated for each of these segments by dividing the number of standing events assigned per segment by the length of the segment. Segments were classified manually as region of no strandings (0 strandings per km), single stranding events (0.01-0.02 strandings per km), low rate of stranding events (0.03-0.08 strandings/km), medium rate of strandings (0.09 to 0.20 strandings/km) and high rate of strandings (>0.21 strandings/km).

#### 2.4.3 Spatio-temporal analysis

##### Space Time Cube

We used space-time cube tool of ArcGIS Pro v. 2.4.2 to integrate spatial and temporal parameters of the data into cubes or bins and store them into a Network Common Data Form (NetCDF) file. We used mid of the month i.e., 15^th^ of each month for the records which did not have precise date information (n =62). These stranding locations were converted into a shape file and then imported into ArcGIS Pro version 2.4.2 to create Space Time Cubes using *Create Space Time Cube Using Aggregating Points* feature to generate patterns. This feature aggregates data points based on chosen spatial and temporal scale (ESRI ArcGIS, 2016) and uses Mann-Kendall statistics to identify trends (Hamed 2009). Mann-Kendall statistics is a rank correlation statistic between ranks of observations and their time sequence. It calculates the values for each bin and identifies the trend based on the z-score and p-value for each bin. A small p-value (0.05) indicates that the trend is statistically significant (Hamed 2009). For this study, the strandings points across 50km were aggregated over 1-year period and further used for emerging hotspot analysis.

##### Emerging Hotspot Analysis

The NetCDF file created in Space-Time Cube tool was taken as an input for “Emerging Hotspot Analysis” tool on ArcGIS Pro which performs Getis-Ord (Gi*) statistics for each bin of space time cube and the neighbouring bins (Manepalli et al. 2011). The Gi* statistics calculates and compares the value of each bin in the space time cube to neighbouring bins and identifies hot spot trends based on the degree of association between two bins (ArcGIS, 2016; Getis and Ord, 1992).

Data was then evaluated into eight types of hot spot and cold spot trends using Mann Kendall statistics. The values of neighbouring bins were also considered while marking any bin as a hot or cold spot.

## 3. RESULTS

We compiled a dataset consisting of 1674 records of marine mammals from 1748 −2017 after removing duplicate reports. It included 660 reports of sightings, 59 reports of induced mortalities or hunting records, 240 reports of incidental mortalities, 632 unique stranding records (live / dead), and 83 records which could not be categorised as either of these because of incomplete information.

### 3.1 General trends

#### 3.1.1 Sightings

A total of 3299 individuals were sighted throughout the Indian coastline between 1748 and 2017 (Figure 1 a). Sighting data on the east coast (species = 18, *n_i_* =1105) was mostly restricted to Odisha and Tamil Nadu (representing 97% of total east coast sightings). Whereas on the west coast, Maharashtra, Gujarat and Karnataka contributed to highest sighting records (representing 85% of total west coast sightings). Sightings from the islands also contributed to 24.85% of the dataset (ANI =24.37 %, LD =0.48 %). Highest incidence of sightings was for dolphins (*n_i_* = 1894) followed by dugongs (*n_i_* = 959) baleen whales (*n_i_* =58) and toothed whales (*n_i_* =17).

**Figure 1 (a):**
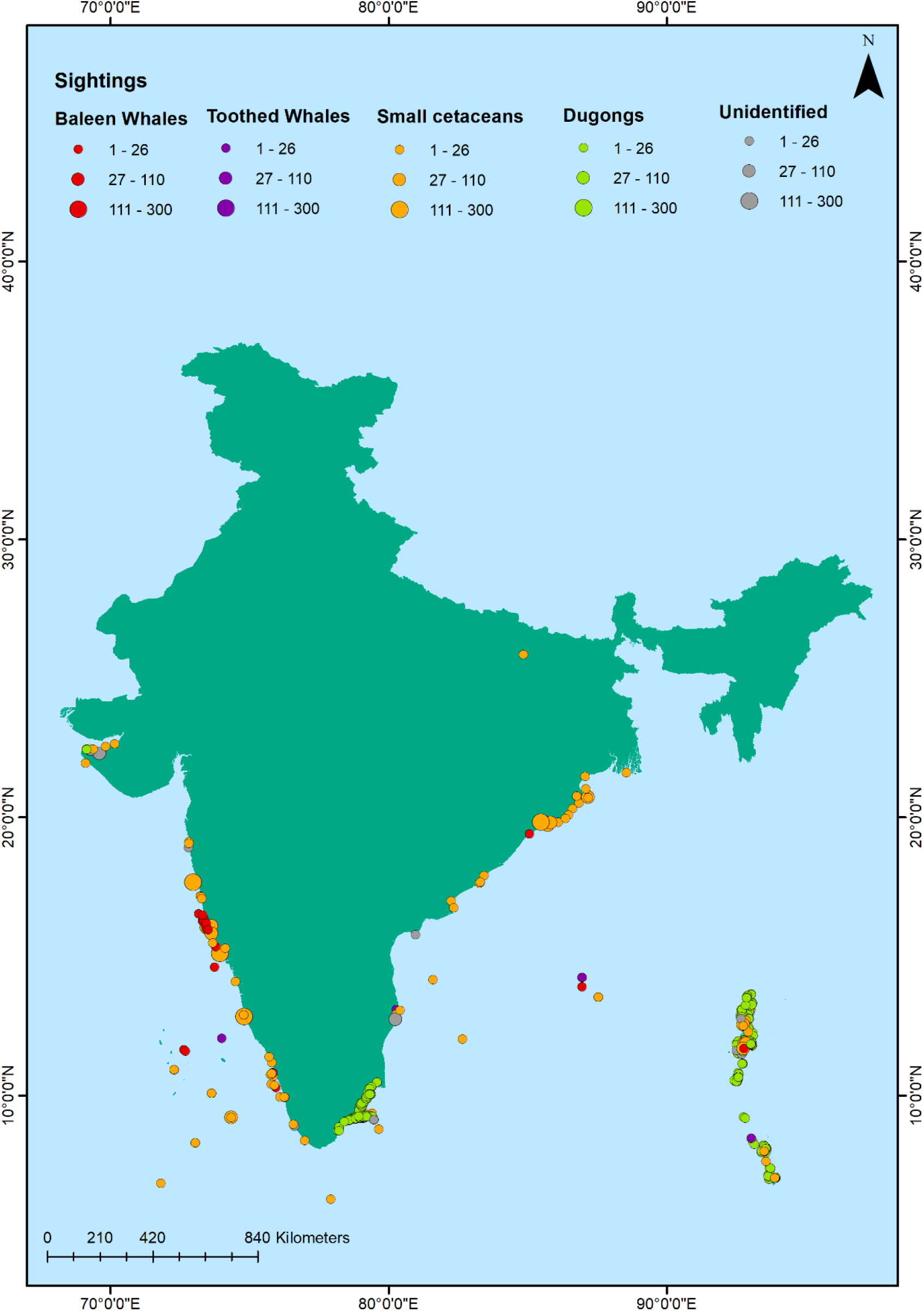
General trends of marine mammal sightings across the Indian coastline.

#### 3.1.2 Induced mortalities

A total of 59 incidences (*n_i_* =102) were recorded of marine mammals being hunted/ captured between the years 1748-2017 (Figure 1 b). The total number of animals hunted/ captured deliberately is similar along east coast (*n_i_* =33), west coast (*n_i_* =29) and islands (*n_i_* =36). Out of all marine mammal species, 90% of the animals hunted at the east coast were *D. dugon* (*n_i_* =30, all from Tamil Nadu). On the west coast, records of hunting incidences of *Neophocaena phocaenoides* were highest (79% of total records on west coast, *n_i_* =23). In the islands (i.e., Andaman and Nicobar Islands), 94% of the hunting records were of dugongs (*n_i_* =34).

**Figure 1 (b):**
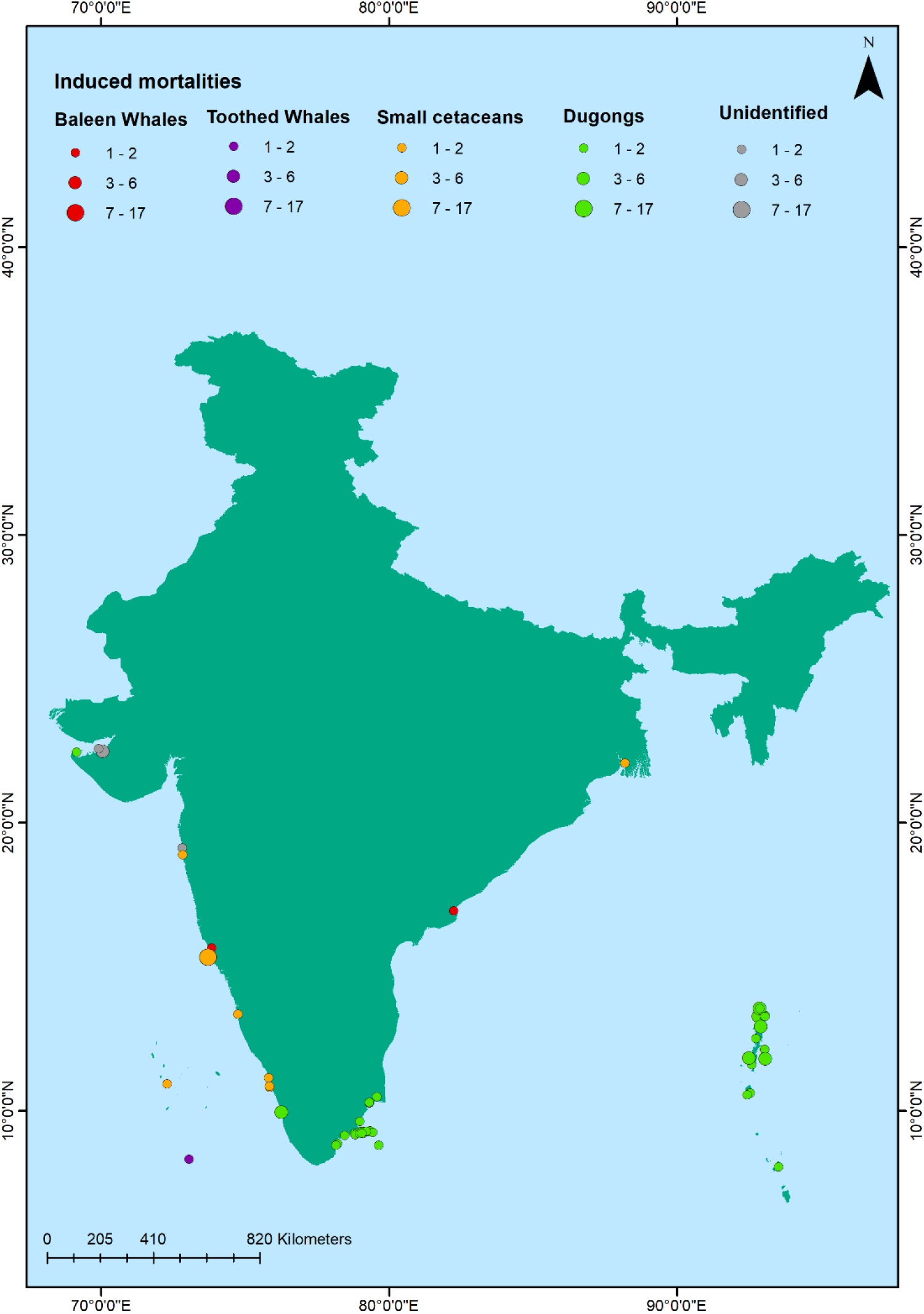
General trends of induced mortalities of marine mammal across the Indian coastline.

#### 3.1.3 Incidental mortalities

A total of 240 accidental net entanglements (*n_i_* = 1356) were reported along the Indian coast between the years 1748 and 2017 (Figure 1 c). The number of animals entangled were similar along east (*n_i_* =670) and west coast (*n_i_* =654) with low reporting from the islands (*n_i_* =26). Similarly, equal number of species were found to be entangled over east and west coast (n=14), whereas in the islands 4 species were recorded in fishnet entanglement. The species composition of entangled animals, however, differs along the coasts of mainland India. *D. dugon* was found to be most frequently entangled along the east coast (63 incidences, *n_i_* = 594, contributing to 56% of the total numbers on east coast), followed by *Tursiops aduncus* (11 incidences, *n_i_* = 14, 9% of the east coast dataset). On the west coast, *T. aduncus* was the most frequently entangled (18 incidences, *n_i_* = 117, contributing to 18% of the west coast dataset), followed by *N. phocaenoides* (17 incidences, *n_i_* = 34, contributing to 17% of the dataset). The total number of small cetaceans being entangled from west coast (*n_i_* =623) were considerably higher than east coast (*n_i_* =68). Whereas, the total number of dugongs being entangled were very high along east coast (i.e., from Tamil Nadu) (*n_i_* =594) as compared to west coast (i.e., Gujarat, *n_i_* =3) and Islands (i.e. Andaman and Nicobar, *n_i_* = 19). *D. dugon* was the most frequently entangled species in the islands (19 incidences, *n_i_* = 19, contributing to 79% of the total numbers in islands dataset) followed by *Pseudorca crassidens* (3 incidences, *n_i_* = 5, contributing to 12% of the islands dataset). Very few baleen or toothed whales were recorded being accidently entangled throughout the Indian coastline (11 incidences, *n_i_* =11).

**Figure 1 (c):**
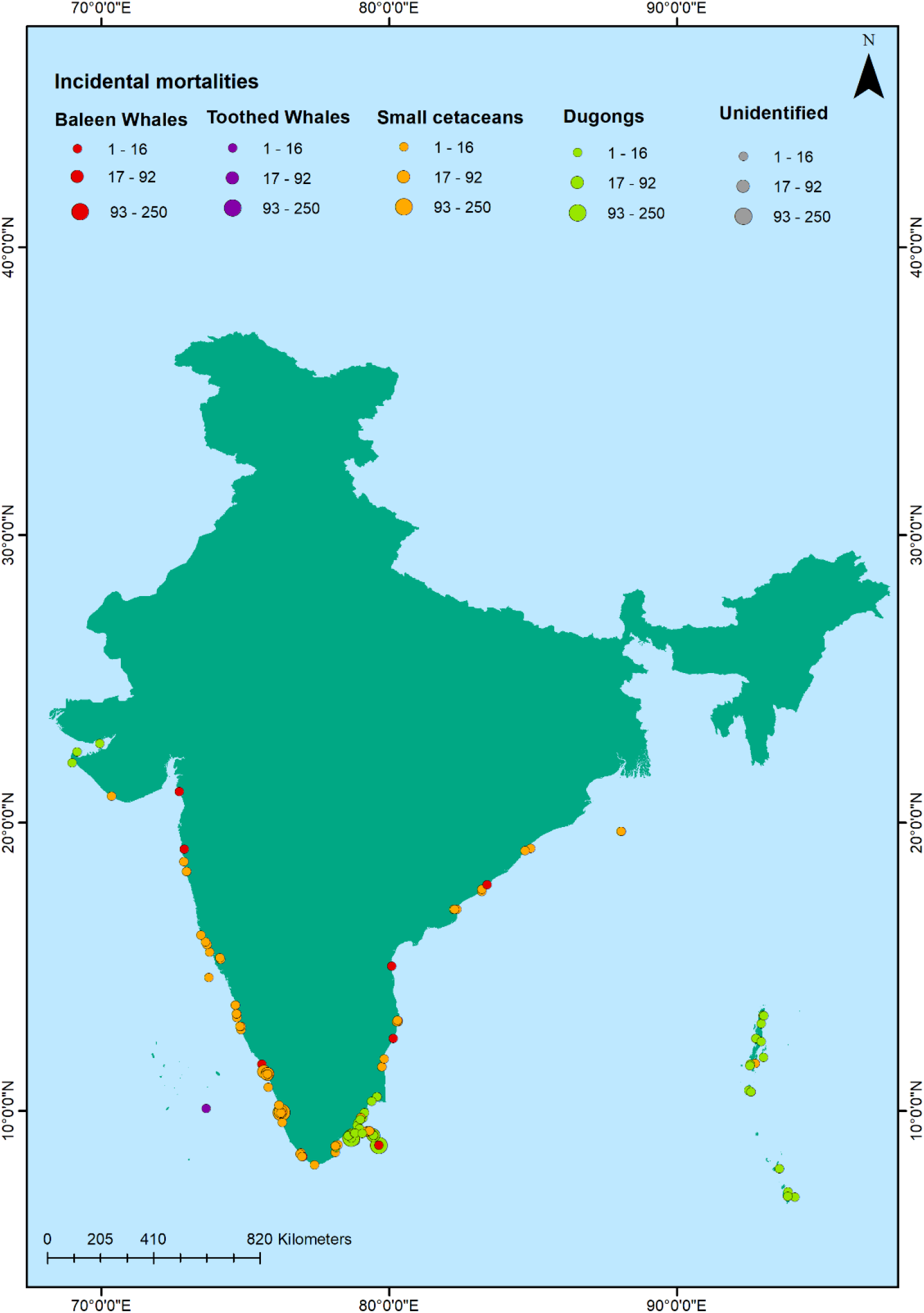
General trends of incidental mortalities of marine mammal across the Indian coastline.

#### 3.1.4 Strandings

Marine mammals stranding reports between the years 1748 and 2017, consisted of ~ 93% dead (*n_i_* = 581) and 7% live strandings (*n_i_* = 51) (Figure 1 d). Considering mass strandings as strandings with *n_i_* > 2 (excluding mother and calf) (D’Amico et al. 2009; Hamilton and Lindsay 2013), 8.5 % of the reports were of mass strandings (21 strandings, *n_i_* = 1054). Most of the records did not have information about the sex of the stranded animal (83%), the age class (88%) or the state of decomposition of the carcass (53%). Out of all stranding incidences, highest number of stranding were reported of dugongs (strandings=190, *n_i_* =228), followed by baleen whales (strandings=178, *n_i_* ==190), smaller cetaceans (strandings=157, *n_i_* ==552) and toothed whales (strandings=47, individuals=48). There were 54 incidences (*n_i_* =54, 9% of total stranding data) where the animal was not identified reliably to include in either of the groups.

**Figure 1 (d):**
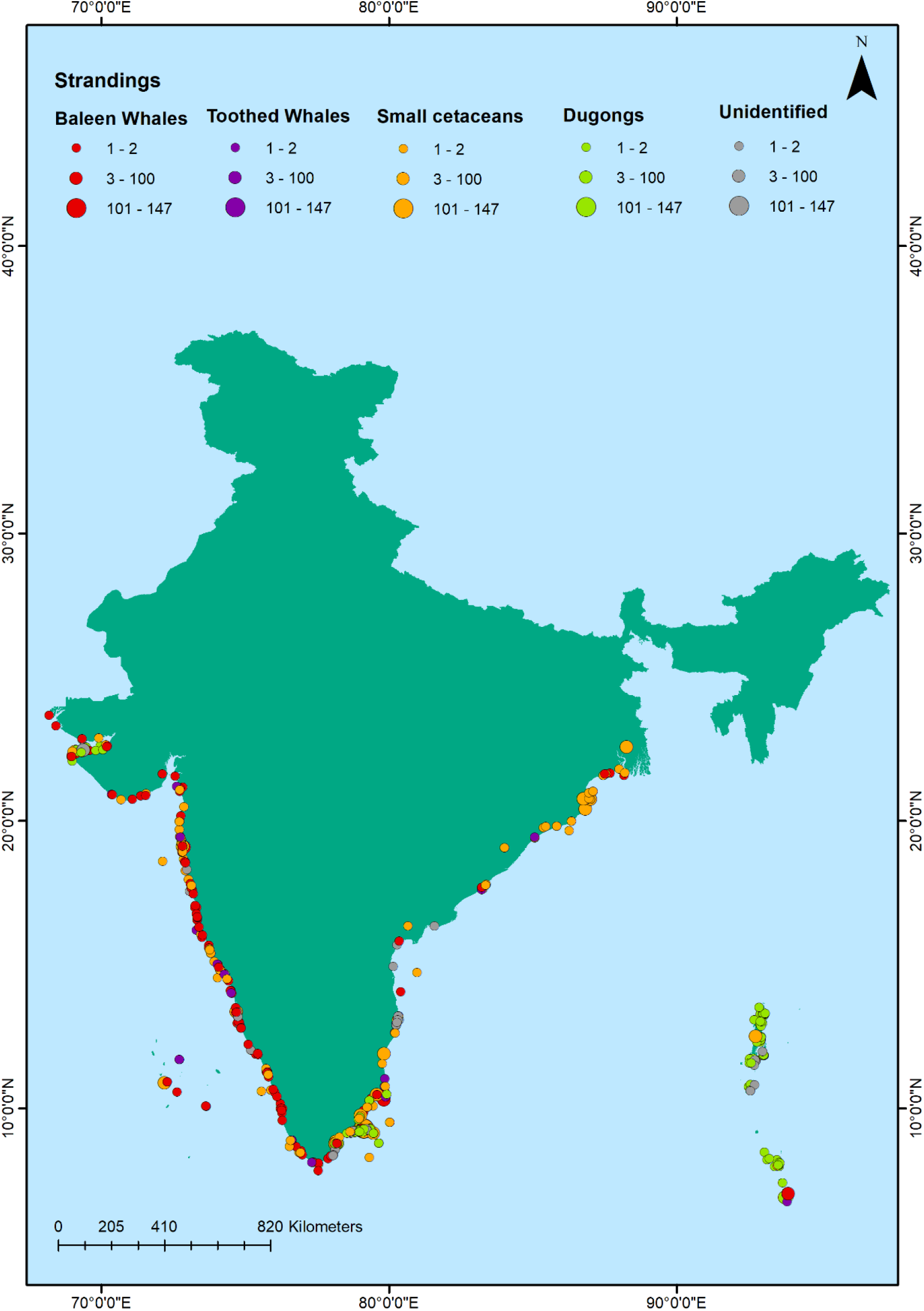
General trends of marine mammal strandings across the Indian coastline.

Species composition and frequencies of strandings were different on east coast, west coast and in the islands. Twenty-two species were found stranded on the east coast with *D. dugon* as the most frequently stranded species (83 incidences, *n_i_* = 107, contributing to 29% of the total numbers on east coast). This was followed by *Sousa chinensis,* (31 incidences, *n_i_* = 108, contributing to 10% of the data). On the west coast, 20 species were recorded as stranded, *Balaenoptera musculus* being the most frequent (28 incidences, *n_i_* = 29, contributing to 12% of the total numbers on west coast) followed by *Neophocaena phocaenoides* (23 incidences, ni = 39, contributing to 10% of the data). In the islands, 13 species were stranded, *Dugong dugon* (93 incidences, *n_i_* = 102, contributing to 77% of the total numbers islands) > *Physeter macrocephalus* (8 incidences, *n_i_* = 8, contributing to 6% of the data; Table 1).

**Table 1:**
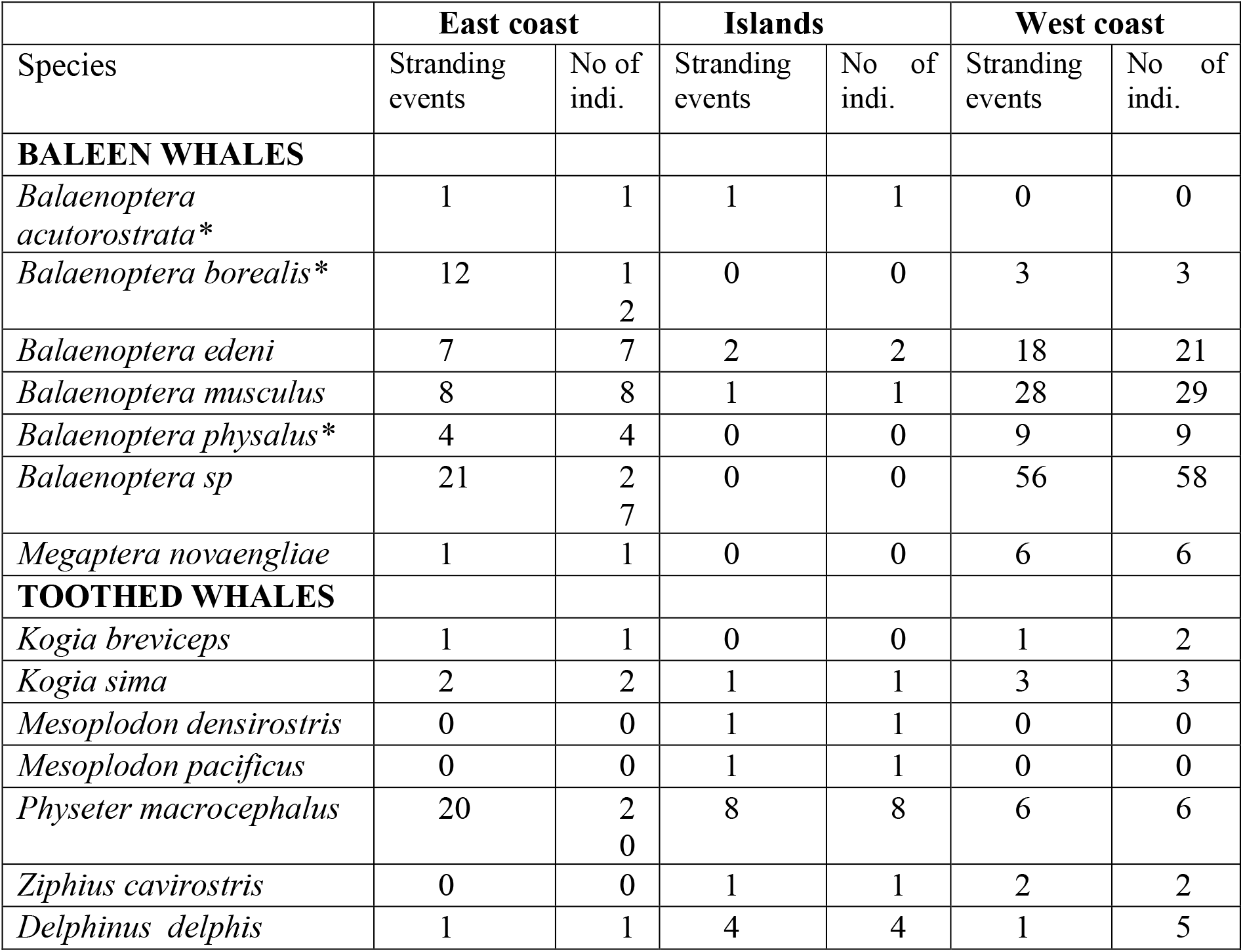
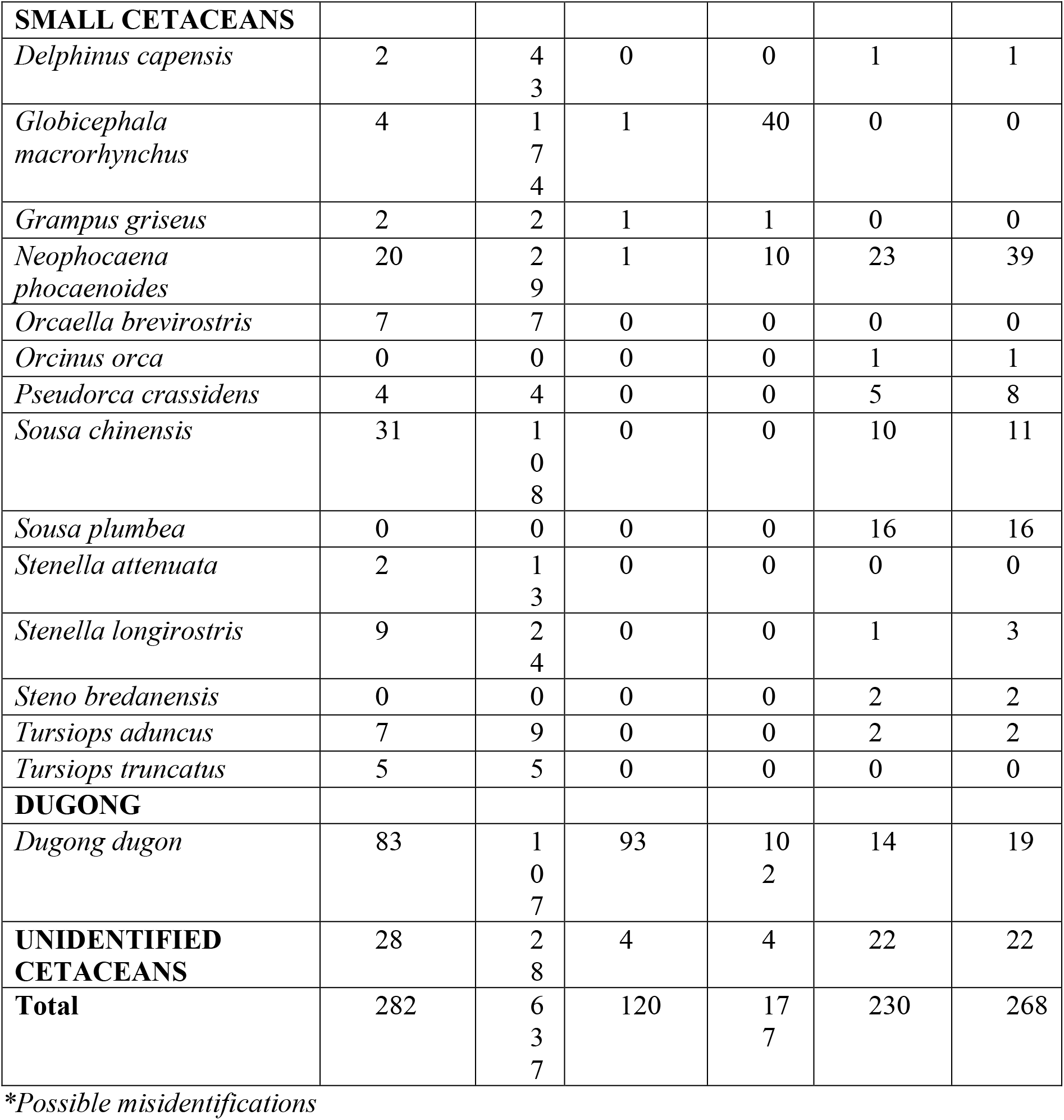
Number of strandings recorded of marine mammal species from east coast, west coast and from islands (Lakshadweep and Andaman & Nicobar Islands) between 1748-2017.

##### a. Baleen whales

Out of the total 178 stranding incidences (*n_i_* =190) of baleen whales, a large number of strandings were unidentified baleen whales (east coast *n_i_*= 27, west coast *n_i_* =58, islands *n_i_* =4; i.e. 47% of the data). Rest of the stranded animals were identified to 6 different species (see Table 1) including misidentified species (no confirmed evidences for *B. acutorostrata, B. borealis and B. physalus* from Indian waters; MMRCNI, 2018). The number of baleen whale strandings were comparatively higher on the west coast (*n_i_* =126), as compared to east coast (*n_i_* =60). The east and west coast received all six species of baleen whales, whereas only three species stranded on the islands. *Balaenoptera borealis* (misidentified) was the most commonly stranded species across the east coast (12 incidences, *n_i_* = 12, contributing to 11% of the data) whereas *B. musculus* was the most frequent across the west coast (28 incidences, *n_i_* = 29, contributing to 11% of the data). Baleen whale strandings were very rare in the islands (4 incidences, *n_i_* = 4). Out of the species identified, *B. edeni* stranded twice (*n_i_* = 2, contributing to 50% of the data).

##### b. Toothed whales

There were 47 strandings (*n_i_* =48) of toothed whales along the Indian coast. The number of strandings of toothed whales were higher on the east coast (*n_i_* =23) as compared to west coast (*n_i_* =13) and the islands (*n_i_* =12). *Physeter macrocephalus* was found to be the most common species to get stranded, and was found throughout the Indian coastline (east coast= *n_i_* = 20, west coast *n_i_* = 6, islands *n_i_* =8; 70% of the data collectively))), whereas *Mesoplodon densirostris* and *Mesoplodon pacificus* were found stranded only once in the islands. *Kogia sima* was found to strand on both the coasts and islands, but in very low numbers (east coast *n_i_*= 2, west coast *n_i_*= 3, islands *n_i_*= 1).

##### c. Smaller cetaceans

There was a total of 157 incidences (*n_i_* =552) of small cetacean strandings belonging to 14 species, out of which 21 were mass strandings. The highest number of individuals in mass stranding event was 147 individuals belonging to species *Globicephala macrorhynchus* at Tuticorin in Tamil Nadu. There were a greater number of smaller cetacean strandings on east coast (*n_i_* =418) as compared to west coast (*n_i_*= 83) and the islands (*n_i_* =51; Table 1). East coast received a higher diversity of stranded small cetaceans (number of species=11) as compared to west coasts (number of species=9) and the islands (number of species=3). *S. chinensis* was the most commonly stranded species across the east coast (31 incidences, *n_i_* = 108, contributing to 33% of the data) whereas *N. phocaenoides* was the most frequent across the west coast (23 incidences, *n_i_* = 39, contributing to 37% of the data; Table 1).

##### d. Dugongs

The current distribution of dugongs in India is in the shallow coastal waters of Gujarat, Tamil Nadu and Andaman & Nicobar Islands (D’ souza and Patankar 2009; Anand et al. 2017). There are 190 stranding events recorded between the years 1893 and 2017. The highest number of stranded animals were recorded from Tamil Nadu (*n_i_* =107) closely followed by Andaman and Nicobar Islands (*n_i_* =102) and very low number of records from Gujarat (*n_i_* =19).

### 3.2. Temporal stranding patterns

Our analysis of temporal trends for the last 42 years (1975-2017) showed that the mean number of strandings along the Indian coast was 11.25 ±9.10/ year. The number of strandings show an increasing trend for two decades after 1975, and then dropped in the decade 1995-2004. The number of stranding events show a distinct rise post 2005 (16.6 ±6.52 / year) with the highest increase being recorded recently (2015-17; 27.66±12.03/year) (Figure 2).

**Figure 2.**
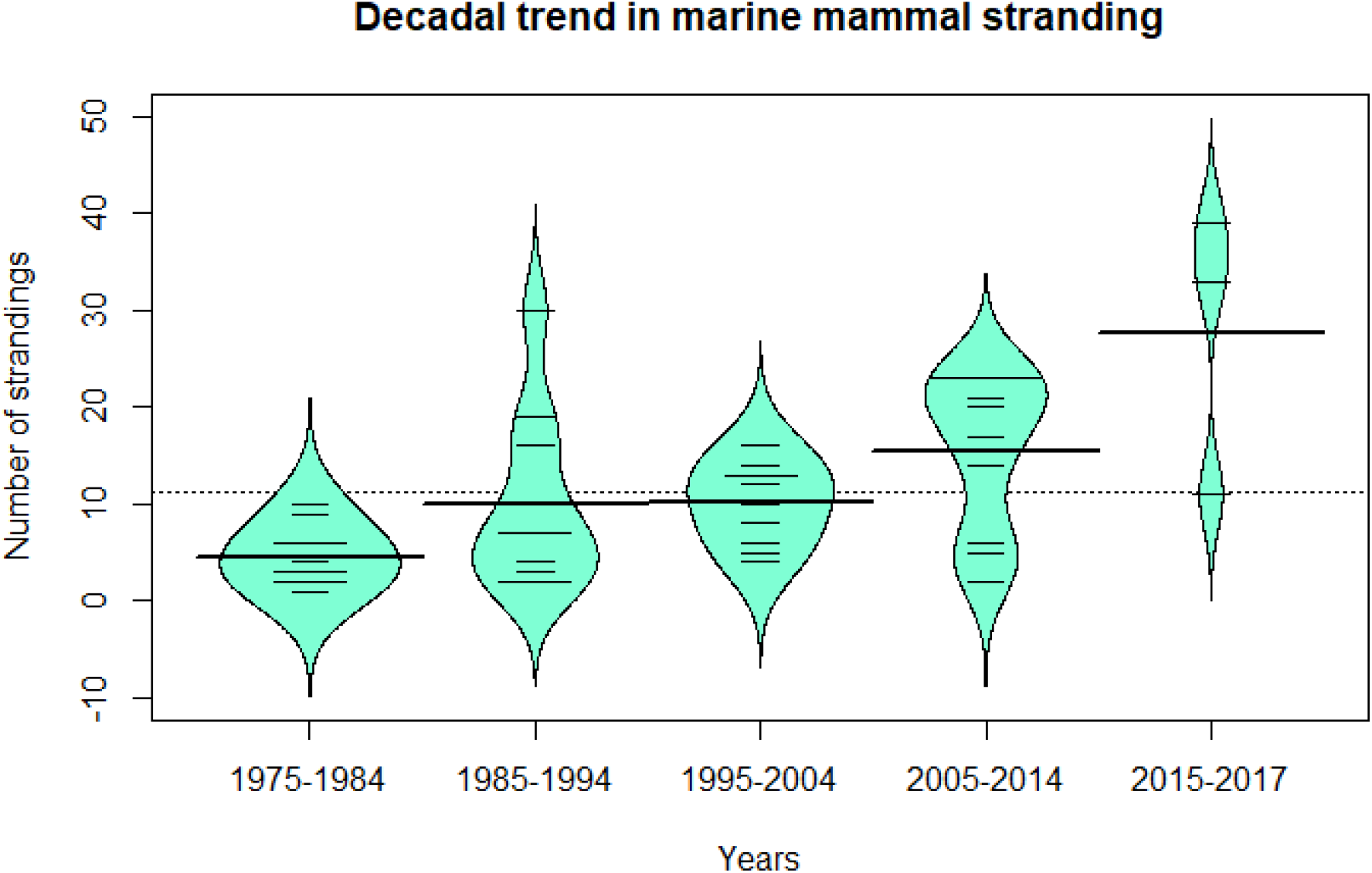
Decadal trends in marine mammal stranding events across the Indian coastline.

#### a. Baleen whales

On west coast, mean stranding rate throughout the years (1975-2017) was 0.0010± 0.0014 strandings/km, and a steady rise was observed in rate of strandings after 2010. A seasonal trend was observed as well, with a peak in the month of September (*s_r_*=0.0061 ±0.0016 strandings/km), i.e. towards the end of monsoon season, and lowest strandings were recorded in the month of June (*s_r_* = 0.0016 ± 0.006 strandings/ km) (Figure 3).

**Figure 3:**
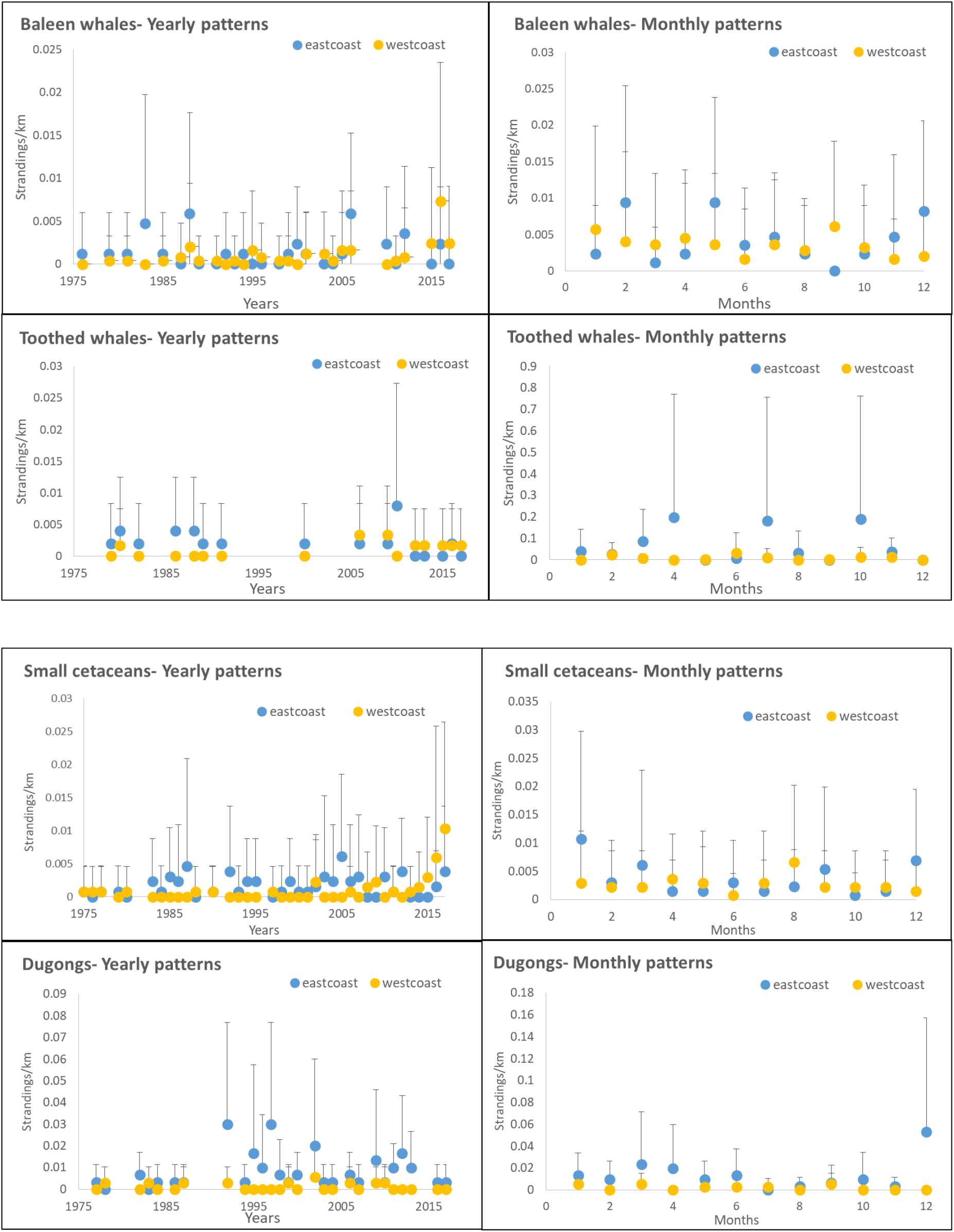
Temporal trends of marine mammal stranding events across the years and months along the Indian coastline.

The mean stranding rate of baleen whales on the east coast through 1975-2017 was 0.0013 ± 0.0017 strandings/km, but no specific trends were observed according to years or seasons. Stranding rates of baleen whales did not differ between east and west coast (Mann-Whitney U test, p value > 0.05).

#### b. Toothed whales

The stranding rates of toothed whales differed significantly across both the coasts (Mann Whitney U test, p value< 0.05). The mean stranding rate of toothed whales on west coast was 0.0010±0.0012 strandings/km, whereas on east coast was 0.0022±0.002 strandings/km. The strandings do not show any specific patterns over the years. The stranding events of toothed whales are too low (n= <5 per month) along both the coasts so no pattern could be deciphered according to seasons as well (Figure 3).

#### c. Small cetaceans

The stranding rates of small cetaceans differed significantly across both the coasts (Mann Whitney U test, p value= 0.00). The mean stranding rates of small cetaceans across the west coast was 0.0009±0.0019 strandings/km, and a definite rise was observed in the rates of strandings after 2014 (Figure 3).

A seasonal trend was observed with a definite rise during monsoon, with highest number of strandings recorded in August (*s_r_* = 0.0066 ±0.013 strandings/ km) (Figure 3). Along the east coast, the mean stranding rate across the years was observed to be 0.0016±0.0015 strandings/km. The number of strandings are seen to increase after November, post retreating monsoon season, with the highest number of strandings in month of January (stranding rate = 0.010 ±0.018 strandings/ km) (Figure 3).

#### d. Dugongs

The stranding rates of dugongs show very high differences across Gujarat (west coast) and Tamil Nadu (east coast) (Mann Whitney U test, p value= <0.0001). The rate of dugong strandings in Gujarat was 0.0010±0.0016 strandings/km, and it was too (n= <5) to comment on seasonal/ monthly patterns. The mean stranding rates through the years on the east coast (i.e. from Tamil Nadu) was 0.0083±0.008 strandings/km and strandings were found to be the highest in the month of December (*s_r_* = 0.0533 ± 0.104strandings/km) (Figure 3).

### 3.3. Spatial patterns

Regions/ coastline near Mumbai (0.38 strandings/km), Kozhikode (0.28 strandings/km), Tuticorin (0.4 strandings/km), Rameshwaram (1.82 strandings/km), Chennai (0.32 strandings/km) and Bhubaneshwar (0.26 strandings/km) were observed to have higher number of stranding events (Figure 4 a). Even though the total number of strandings on east coast were more than double that of west coast (refer Table 1), this analysis reveals that strandings along east coast are concentrated towards Tamil Nadu region rather than being spread out evenly along the coast. On the other hand, strandings on west coast are more or less evenly spread out.

**Figure 4 (a):**
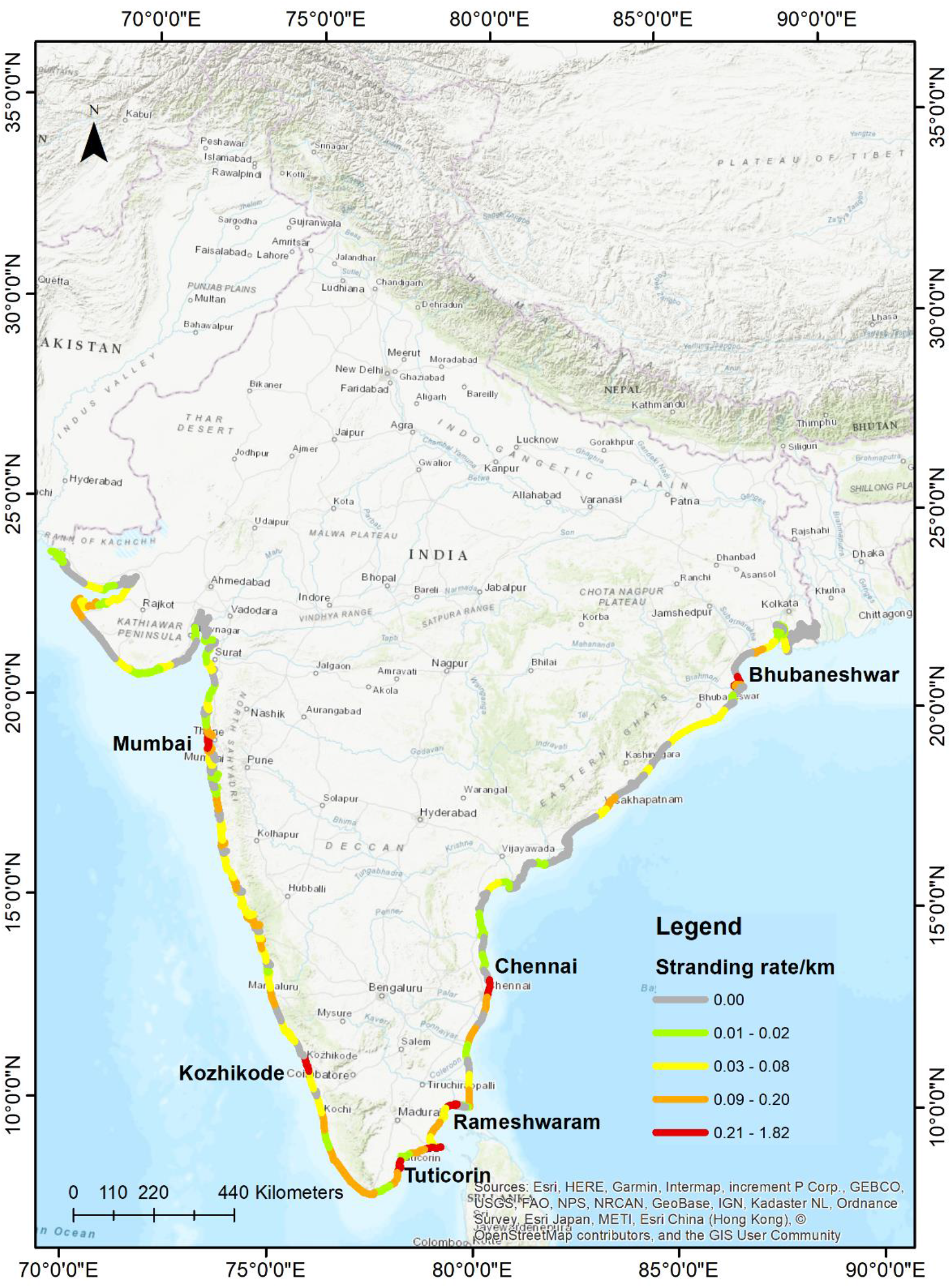
Rate of strandings (number of strandings/ km) of marine mammals along the Indian coastline.

#### 3.3.1 Emerging Hotspot Analysis

Only 434 out of the total 632 strandings could be used for this as rest of the records did not have information about month of stranding. The analysis detected four hotspot categories: no patterns, consecutive, sporadic and intensifying hotspots (see Table 2).

**Table 2:**
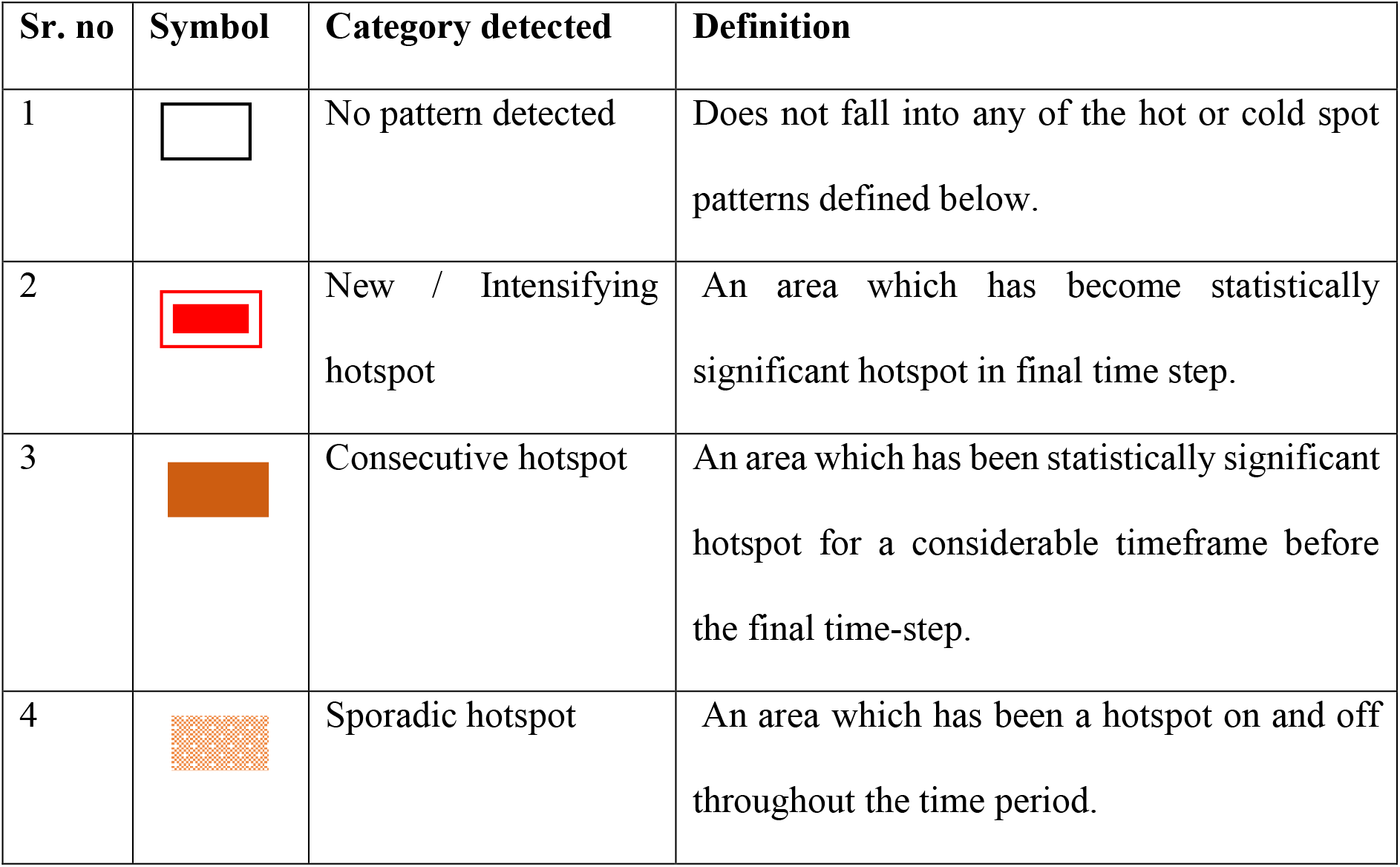
Emerging hotspot categories detected and their definitions

| Sr. no | Symbol | Category detected | Definition |
| --- | --- | --- | --- |
| 1 |  | No pattern detected | Does not fall into any of the hot or cold spot patterns defined below. |
| 2 |  | New / Intensifying hotspot | An area which has become statistically significant hotspot in final time step. |
| 3 |  | Consecutive hotspot | An area which has been statistically significant hotspot for a considerable timeframe before the final time-step. |
| 4 |  | Sporadic hotspot | An area which has been a hotspot on and off throughout the time period. |

Along the west coast, the southern region of Gujarat, near Veraval (district Gir Somnath) and the coast of Surat emerged as new/intensifying hotspots (Figure 4b). It implies that these regions were never a hotspot but the frequent strandings in the last time step, i.e., in 2017, being statistically significant, highlight them as new/ intensifying hotspots.

**Figure 4 (b):**
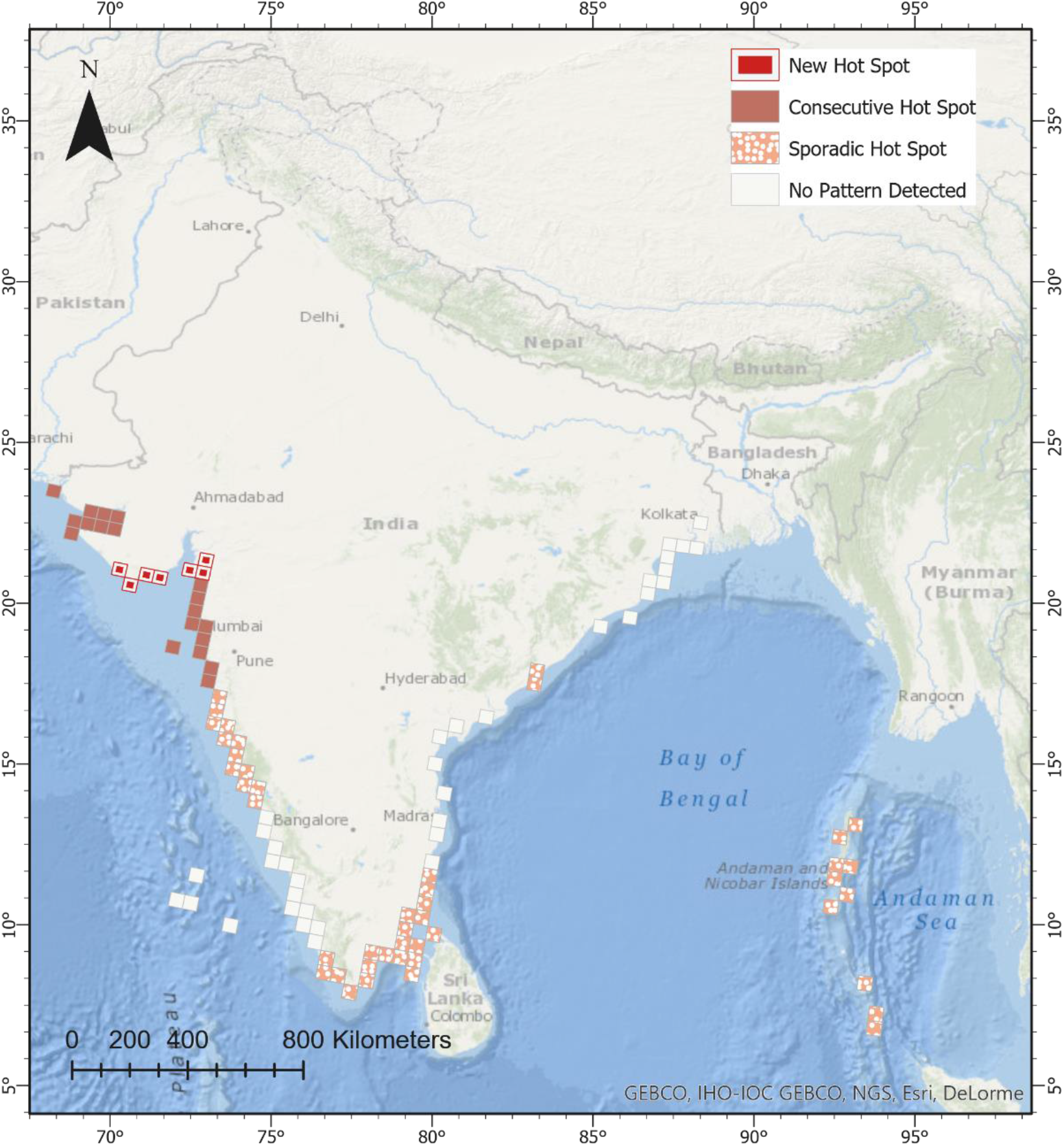
Emerging hotspots for all stranded marine mammals across the Indian coastline.

The area around Gulf of Kutch Marine National Park and most of the Konkan coast, except Ratnagiri, have not been significant for marine mammal strandings in the earlier years, but the strandings in few recent years have made this part statistically significant resulting in consecutive hotspot pattern (less than 90% bins being statistically significant). Further, the northern Karnataka coast, south of Kerala and Tamil Nadu, Vishakhapatnam and Andaman and Nicobar Islands are sporadic hotspots. These regions did receive significant number of strandings, but they were inconsistent throughout the time period (ESRI, ArcGIS Pro, 2016) thus making it difficult to be demarcated as hotspots. Further research is needed to ascertain any pattern in these areas. No significant pattern was detected at southern Karnataka, northern Kerala, Lakshadweep Islands, Pondicherry, Andhra Pradesh, Odisha and West Bengal coast.

##### a. Baleen whales

The baleen whale strandings resulted in three patterns. The Gujarat and Maharashtra coasts are consecutive hotspots, which means baleen whale strandings in recent few years (i.e. 2015-2017) have made this region a hotspot. Northern part of Karnataka and Kanyakumari are sporadic hotspots, which means they have been hotspots for baleen whale strandings on and off throughout the study period (Figure 4 c). No pattern was detected from rest of the strandings of baleen whales along the coast.

**Figure 4 (c):**
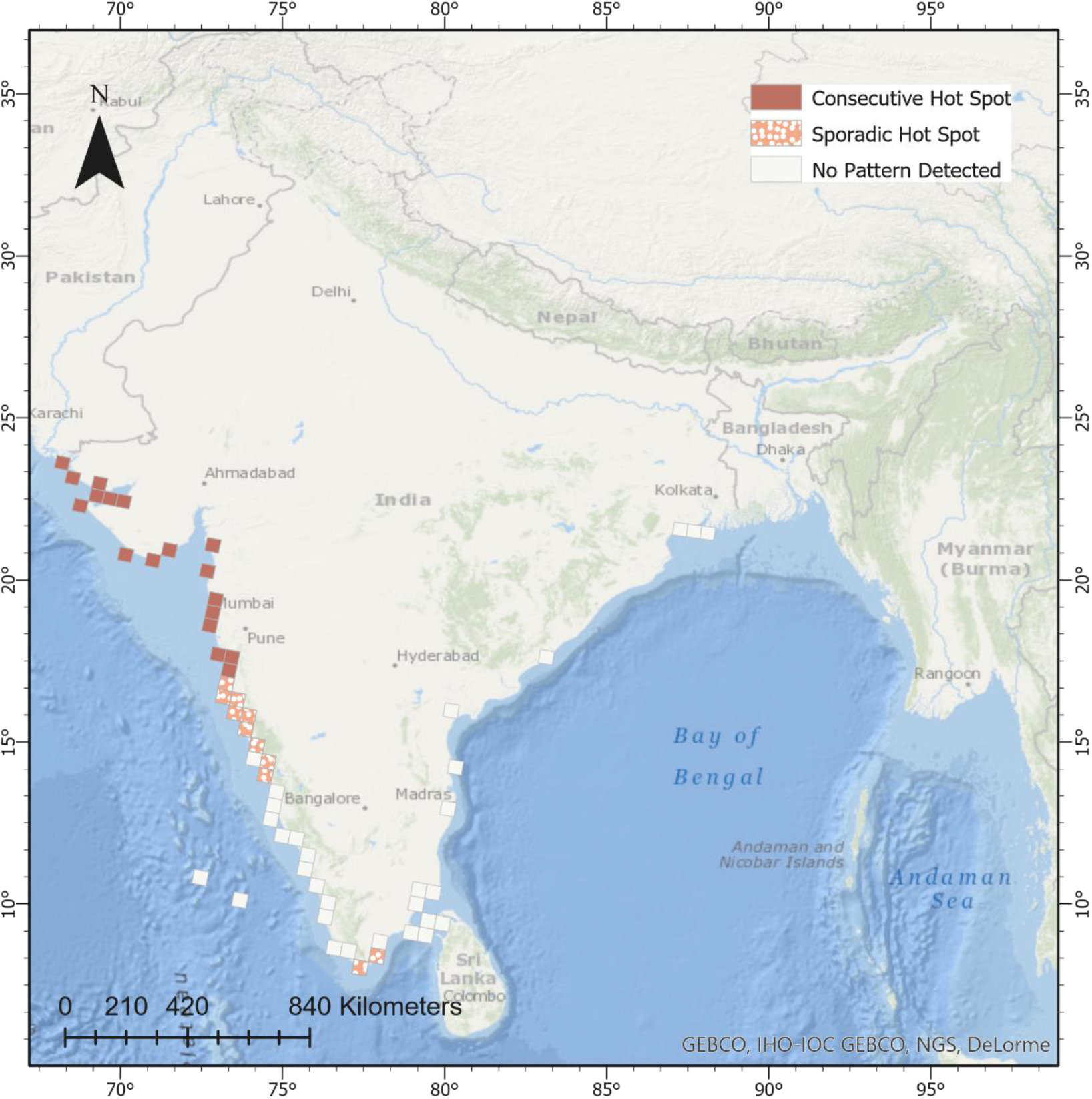
Emerging hotspots for all stranded baleen whales across the Indian coastline.

##### b. Small cetaceans

The strandings of small cetaceans-dolphins and finless porpoises have been sporadic over most of the coast, except the region between Mumbai and Ratnagiri, which is a consecutive hotspot. No patterns were detected from strandings along north of Andhra Pradesh, Odisha, West Bengal, Lakshadweep and Andaman Islands (Figure 4 d). As this tool required a certain minimum number of data points to analyse, i.e. 60, the stranding records of toothed whales and dugongs were too low to process (ESRI, ArcGIS Pro version 2.4.2).

**Figure 4 (d):**
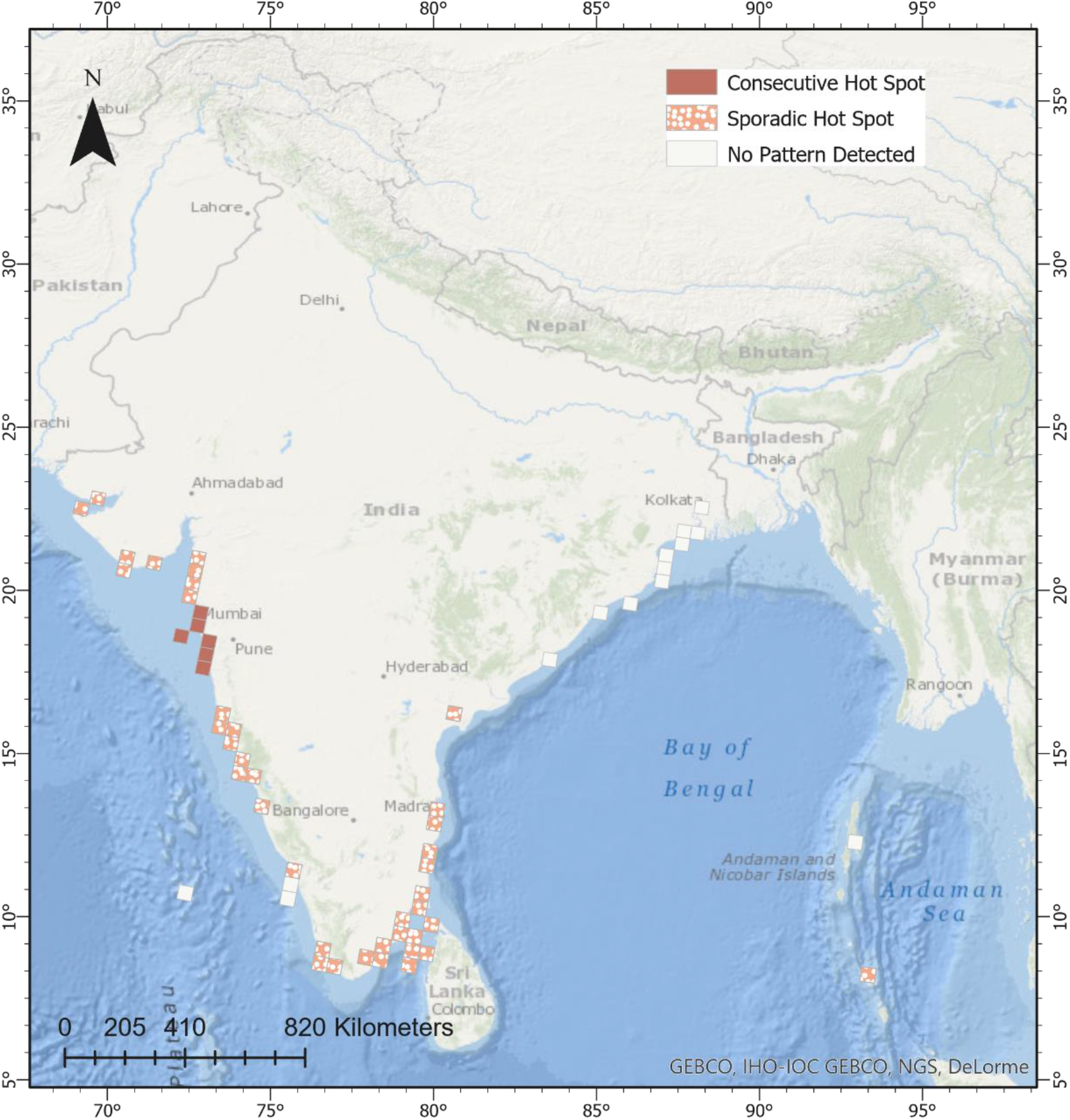
Emerging hotspots for all stranded small cetaceans across the Indian coastline.

## 4. DISCUSSION

This study is the first attempt to utilise publicly available data on marine mammal strandings to identify stranding hotspots in India. The dataset used in the study was compiled from scientifically vetted databases, primary surveys, government reports and newspaper articles providing a comprehensive synthesis of long-term marine mammal stranding records in the country. Even though temporally discontinuous and lacking uniformity in reporting parameters, this dataset provides critical information for managing marine mammal strandings across the vast Indian coastline. Adding over 752 unique records to the existing MMRCNI database, this dataset helps us a) to highlight general patterns in marine mammal occurrence b) provide evidence for emerging stranding hotspots and c) delineate areas for immediate establishment of a robust stranding response program along the Indian coastline. We highlight the importance of using a novel GIS tool to identify patterns in the marine mammal strandings over an extensive coastline. Finally, we identify regions of high priority to monitor strandings and provide recommendations for developing a robust stranding response mechanism in the country.

Reports of marine mammal occurrence have steadily increased over the last two decades reflective of increased survey effort and improved awareness levels, necessitating a systematic investigation. Sightings records on the east and west coast differ considerably due to differences in efforts undertaken to survey marine mammals. Maximum sighting reports were of dugongs obtained through dedicated interview surveys at its distribution sites (Sivakumar & Nair, 2013) followed by those of Indo-pacific humpback dolphins through the studies conducted on the west coast (Sutaria and Jefferson 2004; Sutaria et al. 2015b; Bopardikar et al. 2018). Hunting or capture data identified finless porpoise on the west coast (from the years 1827-1983) and dugongs on the east coast being hunted in the past as well as from records in the past ~100 years (records from 1910-2013). For dugongs, the dataset points towards specific areas in Tamil Nadu (Tuticorin and Ramanathapuram districts) where extensive monitoring is required. Identical patterns in fishnet entanglement appear from both coasts with dugong being most frequently entangled on the Tamil Nadu coast, small cetaceans being reported widely from all the west coast with very few reports of baleen or toothed whale entanglements. Most stranding events comprise of single individuals with little to no information on the condition of the carcass, gender identification or age class. Further, over 8% stranded individuals could not be identified till species level with several misidentifications in the dataset. High dugong strandings have been reported through focused surveys (Sivakumar and Nair 2013) and formation of volunteer networks (Hatkar et al. 2020; Srinivas et al. 2020) but these records come from isolated pockets of dugong distribution along the Indian coast. This absence of quality information collection on stranding events arises due to lack of appropriate facilities and trained manpower to report strandings and conduct necropsies throughout the Indian coastline. Further, most biological information is lost with a delay in reporting and/or misidentification at the stranding site requiring strong coordination with government agencies and the scientific community for an appropriate stranding response (Kumarran, 2012).

Marine mammals migrate locally (Dong et al. 2017) as well as long-distance (Horton et al. 2011) in response to changes in habitat characteristics (such as sea surface temperature or Harmful Algal Blooms), to breed or to find optimal foraging regions (Boyd 2004; Truchon et al. 2013). Long-term stranding datasets facilitate monitoring these movements, identifying novel breeding, or foraging habitats and decode linkages between climate variability and marine mammal communities along the coastline (Truchon et al., 2013). We observed a distinct rise in strandings over decadal time scales post 2005, with the highest frequency of strandings noted between 2015-17 (ns/year = 27.66, 13% of total strandings). This rate is arguably the highest stranding rates recorded from any region of the world (Segawa and Kemper 2015; Alvarado-Rybak et al. 2020). Among groups, baleen whale strandings show a steady increase along west coast post 2010, more reports coming from September (i.e., post-monsoon) while lowest were reported in June. No specific seasonal trends of baleen whales were observed along east coast. Toothed whales did not show any specific seasonal patterns with very few records from either coast. Small cetaceans show an increase in strandings after 2014 along the west coast esp. during August (i.e., post-monsoon). Along east coast, the small cetacean strandings rise after November peaking in January. For dugongs, numbers along west coast were too low to detect any seasonal patterns but along east coast (Tamil Nadu), the strandings peak in December. These patterns emerge from a multi-scale (coasts) and multi-species dataset illustrating the complexity, seasonality, varying anthropogenic pressure and reporting differences of the stranding events. Suitable management interventions and focused monitoring is required along coastal sections where higher strandings have been reported in these specific seasons to decipher the precise causes.

Globally, spatial patterns of marine mammal strandings point towards localised threats in certain regions e.g. use of high powered SONARs for bathymetric studies or for military use (Firestone and Jarvis 2007; Lurton and DeRuiter 2011), deep sea oil explorations (Smodlaka et al. 2019), negative interactions with fishing gears (Ahn 2017), or vessel strikes near shipping ports or high tourism areas (Groom et al. 2004). Our spatial analysis revealed that the geographical distribution of the strandings was not homogenous and division of the coastline into sections revealed the patterns on a finer scale (50 Km). We found that the stranding rates on the sections of west coast were oscillating between low to moderate, with a few sections (districts of Mumbai and Kozhikode) displaying high stranding rates. On the contrary, the southern sections of the east coast had low to moderate strandings with higher rates from Tamil Nadu, whereas the coastline in the central and northern region had negligible stranding rates, with one or two exceptions.

Spatial stranding patterns of marine mammals in India coincide with monsoon winds which flow towards the landmass at the respective coast (Shankar 2000; Kripalani and Kumar 2004). The monsoon season differs along both coasts as south-west winds bring rains to west coast in the months of June-September, whereas the east coast experiences retreating monsoonal winds from the north-eastern direction, in the months of October-November (Gadgil, 2003). This changes the ocean currents and sea surface temperature patterns during two distinct periods across the Indian subcontinent (Shankar 2000; Kripalani and Kumar 2004). Post-mortem deposition of carcasses along certain sections of the coastline are thus governed by these environmental factors (Norman et al. 2004; Leeney et al. 2008a; Truchon et al. 2013). The observed patterns are thus possibly be an artifact of seasonal changes in these hydrodynamic drivers and are likely to increase stranding events when the currents drive the animals towards the shore, or reduce if the dead animals are driven away into deeper regions of the ocean. These results highlight the need of validated reporting of strandings along the Indian coastline to understand the effect of climatic drivers which are likely to be highly informative over longer time frame (Tomás et al. 2008; Sepúlveda et al. 2020).

Increased stranding rates might relate to extreme climatic conditions (Meager and Limpus 2014; Sepúlveda et al., 2020), oceanographic anomalies (Hart et al. 2006), exposure to pollutants (Hall et al. 2006) and other biological events (e.g. red tides, diseases) (Flewelling et al. 2005; Gulland and Hall 2007; de la Riva et al. 2009). Besides these, certain human activities such as oil spills (Wilkin et al. 2017), oil exploration, increased intensity of fishing (Leeney et al. 2008), illegal fishing practices (Aragones et al. 2010) and high-speed recreational vessel traffic (Groom et al. 2004) pose localised threats affecting strandings. We cannot deny the possibility that these events might result in different spatial patterns of strandings across finer or greater spatial scales than the one chosen for the current study. However, the patterns obtained in this study provide a baseline to undertake investigations considering data on the species distribution, localised threats, and variations in physical oceanographic processes.

Despite the challenges and limitations of working with an opportunistic database, our study has revealed strong spatio-temporal patterns using the novel ‘Space Time Mining’ tool which offers significant benefits to both conservation biologists and management agencies. It is evident from the Emerging Hotspot Analysis that regions with higher rate of strandings show heterogeneity in stranding frequencies. South-eastern coast shows sporadic nature of stranding events despite having multiple sections having moderate to high stranding rates. On the other hand, consecutive and intensifying hotspots emerging in the north western region draw attention towards determining the causes of a steady increase in stranding rates.

Over 33% of the coastal districts have reported < 5 strandings over the last ~270 years (maximum 7 districts in Gujarat followed by 5 in Kerala) indicating a lack of effort or reporting mechanism in the region. With data limitation arising from reporting bias, the patterns in the study emerge from a temporally disjunct dataset coming from various sources lacking homogeneity in the way information was obtained. The data was collected on an opportunistic basis rather than dedicated beach surveys, and thus the patterns obtained could be biased by factors which are likely to influence the behaviour of informants, such as the accessibility of the shore and weather conditions (Evans and Hammond 2004; Olson et al. 2020) and unequal sampling effort (Huggins et al. 2015). In the Indian context, differences in data availability also arises with kind of marine mammal research undertaken, such as by fishery biologists in the past (Kumaran 2002; Kumarran 2012) with focus in regions in proximity of the fishery institutions (such as Central Marine Fishery Research Institute), or other institutions (National Institute of Oceanography) or individual species or area focused efforts (e.g. along south Maharashtra coast - Konkan Cetacean Research Team; for Indo-pacific humpback dolphins - (Sutaria and Jefferson 2004; Sutaria et al. 2015; Bopardikar et al. 2018; Jog et al. 2018). This is likely to influence the proportion of strandings of that species which dominate the dataset, and patterns might not be generalised or even applicable to the whole group as considered in the study, due to differences in their biology and behaviour. Similarly, the data received from the residents and fishermen of any region is a derivation of the enthusiasm and level of awareness regarding the importance of reporting such events on the public platforms like the media and online databases, or to the researchers working on marine mammals in the area. It also cannot be denied that the sections of the coastlines with higher rate of strandings or which show consecutive or intensifying hotspots are subject to higher level of awareness about marine mammals, have proximity to oceanography/ fishery institutions or have focused research carried out for a certain marine mammal species.

Overall, our results highlight critical areas to be prioritized for monitoring marine mammal strandings in the country. As timely response and documentation is of utmost importance while dealing with stranding events, we recommend setting up of regional marine mammal stranding response centres at the identified hotspots coordinated by a National Stranding Centre. These stranding response centres should consist of a dedicated team comprising of trained personnel and veterinarians from State Forest Department, State Fisheries Department with a network of volunteers from scientific community and general public. This network could be established in line with the recommendations in the 3^rd^ National Wildlife Action Plan of India (https://wii.gov.in/nwap_2017_31) where a National Marine Animal Research and Response Centre with state-wise stranding centres has been proposed. Adequate funding should be made available to facilitate periodic training of manpower and recruitment of scientific personnel at these centres from the centrally-sponsored schemes. Periodic stranding response programs for training field veterinarians, frontline personnel (especially around stranding hotspots) need to be undertaken through central or state sponsored programs. Standard stranding response protocols have proven to increase the efficiency of responding and recording of stranding events (Aragones et al. 2010; Barbieri et al. 2013), and such measures need to be adopted at the earliest. Though data collated at the national level is a challenging task, setting up of a National Stranding Database (as envisaged in the draft National Framework and Guidelines for The Management of Marine Animal Stranding in India) with involvement of various stakeholders would help develop a comprehensive picture of marine mammal populations in Indian waters.

## ACKNOWLEDGEMENTS

We thank CAMPA Authority of the Ministry of Environment, Forest and Climate Change, Government of India for funding this study and members of ‘Recovery of Dugongs and their habitats in India’ team for the help in compilation of the database. Special thanks to Sagar, Sameeha, Rukmini and Sumit for helping in data collation. We are grateful to Director and Dean, Wildlife Institute of India for their constant support to this study. We would also like to thank Marine Mammal Research and Conservation Network of India, State Forest Departments of Gujarat, Andaman and Nicobar and Tamil Nadu for their contribution to the stranding dataset.

## Notes

All authors have approved the manuscript and declare no conflict of interest.

### Competing Interest Statement

The authors have declared no competing interest.

